# Rapid and unbiased enrichment of extracellular vesicles via meticulously engineered peptide

**DOI:** 10.1101/2023.08.04.551951

**Authors:** Le Wang, Zhou Gong, Ming Wang, Yi-Zhong Liang, Jing Zhao, Qi Xie, Xiao-Wei Wu, Qin-Ying Li, Cong Zhang, Li-Yun Ma, Si-Yang Zheng, Ming Jiang, Xu Yu, Li Xu

**Affiliations:** Tongji School of Pharmacy, Huazhong University of Science and Technology, Wuhan, 430030, China; State Key Laboratory of Magnetic Resonance and Atomic Molecular Physics, Innovation Academy for Precision Measurement Science and Technology Chinese Academy of Sciences, Wuhan, Hubei 430071, China; Department of Clinical Laboratory, Renmin Hospital of Wuhan University, Wuhan, 430060, China; Department of Oncology, Tongji Hospital, Tongji Medical College, Huazhong University of Science and Technology, Wuhan 430030, China; College of Horticulture and Forestry Sciences, Huazhong Agricultural University, Wuhan 430070, China; Department of Thoracic Surgery, Tongji Hospital, Tongji Medical Collage of Huazhong University of Science and Technology, Wuhan 430030, China; Department of Pharmacy, Union Hospital, Tongji Medical College, Huazhong University of Science and Technology, Wuhan 430022, China; Department of Electrical Engineering and Department of Biomedical Engineering, Carnegie Mellon University, Pittsburgh, Pennsylvania 15213, United States; Hubei Jiangxia Laboratory, Wuhan 430200, China

**Keywords:** extracellular vesicles, peptide, protein assay, DNA mutation detection, functionalized interface

## Abstract

Extracellular vesicles (EVs) have garnered significant attention in biomedical applications, particularly as biomarkers and therapeutic agents for cancer diagnosis and treatment. However, the rapid, efficient, and unbiased separation of EVs from complex biological fluids remains a challenge due to their heterogeneity and low abundance concentration in biofluids. Herein, we report a novel approach to reconfigure and modify an artificial insertion peptide for the rapid isolation of EVs in 20 min with ∼ 80% recovery. By inserting the peptide into the phospholipid bilayer of EVs, our method enables the unbiased isolation of EVs. Moreover, our approach demonstrates exceptional anti-interference capability and achieves a high purity of EVs comparable to standard ultracentrifugation and other methods. Importantly, we show that the isolated EVs could be directly applied for downstream protein and nucleic acids analyses, including proteomics analysis, exome sequencing analysis, as well as the detection of EGFR and KRAS gene mutation in clinical plasma samples. Our approach offers new possibilities for utilizing EVs in cancer diagnostics through liquid biopsy, as well as in various other biomedical applications.

## Introduction

Extracellular vesicles (EVs) are cell secreted vesicle substances with phospholipid bilayer structure and their sizes are ranging from 30 nm to 1000 nm^1–3^. EVs are present in various body fluids, such as blood, urine, tear ascites and saliva. Previous studies reveal that EVs contain diverse nucleic acids (RNA and DNA), proteins, lipid and other substances inheriting from their original parent cells and participating in various physiological activities, such as intercellular communication^4^, tissue repair and regeneration^5^, and immunomodulation^6^. Additionally, numerous evidences have shown that EVs play important roles in tumor progression processes, including epithelial-mesenchymal transition^7^, angiogenesis^8^, immune system avoidance^9^, cancer migration, invasion and metastasis^10, 11^, and drug resistance^12^. Thus, EVs have been recognized as promising biomarkers for liquid biopsy, which is crucial for early diagnosis of cancers and improvement of cancer survival rate^11, 13, 14^. However, the detection of tumor-associated EVs encounters challenges due to their ultra-trace concentration at the early stage of cancer and the complex matrix interference in the sample^4, 15, 16^. Furthermore, the purity of the isolated EVs might be affected by the co-existence of other biological interference, such as highly abundant lipoproteins and circulating cell-free nucleic acids. Therefore, the rapid, simple and highly efficient isolation and enrichment of EVs from complex clinical samples is the first step and key issue for the EVs based biomedical applications^17–20^.

Several label-free methods have been developed for the isolation and enrichment of EVs, including the gold standard method of ultracentrifugation (UC)^21, 22^, polymer-based precipitation^23^ and microfluidic-based methods (e.g., acoustofluidics, dielectrophoretic separation, and deterministic lateral displacement)^24–26^. However, the UC method for enriching EVs often requires large sample volumes, multiple tedious treatment steps, and a relatively long processing time (∼ 4 h), resulting in poor yields of EVs (5% - 20%)^21^. Polymer-based precipitation methods, such as the polyethylene glycol (PEG) based precipitation, can obtain a relatively high amount of EVs, but they sometimes cause serious aggregation and provide unacceptable purity. Although microfluidic-based label-free methods for EVs enrichment show great promise, the cumbersome chip fabrication and operation procedures, time-consuming process, and the need of professional staff limit their wide applications in clinical trials^21, 27–30^. Except of those approaches, immunoaffinity approaches, such as specific antibody or aptamers-based isolation methods, have also been applied for the separation of EVs^18^ ^31, 32^. However, the EVs exhibit strong heterogeneity^33, 34^, which could seriously impede the efficient enrichment of tumor-associated EVs through immunoaffinity approaches, attributed to the low or lack of relevant antigen expression on heterogenous EVs^4, 33, 35^. This may lead to the omission of crucial cancer-related information, due to the low abundance of the tumor-associated EVs in the early stage of the cancer. Meanwhile, the heterogeneity may also hinder the discovery of new biomarkers on EVs for cancer diagnosis^4^. Moreover, the high cost and challenging storage of the antibodies, as well as the susceptibility to degradation of the aptamers, have also hindered the efficiency of EVs’ capture. To address the aforementioned challenge, we previously used a lipid-nanoprobe (LNP), biotin-tagged 1,2-distearoyl-sn-glycero-3-phosphethanolamine-poly(ethylene glycol) (DSPE-PEG), inserted into the membranes of EVs for the separation and enrichment of EVs^36^. The results demonstrated that more than 80% of EVs were successfully recovered in the buffer solution. However, in clinical plasma samples, only ∼ 50% capture efficiency was achieved, which might be attributed to the interference of complex ingredients in plasma. With the similar idea, a recent study reported efficient and unbiased isolation of EVs by using choline phosphate-functionalized magnetic beads^37^. The isolation mechanism relied on the coordination between choline phosphate on the magnetic beads and phosphatidylcholine on the EV membranes. The captured EVs could be released by simply elevating the temperature of the solution. Although this study was impressive, the process of functionalizing magnetic beads with choline phosphate was complicated. Thus, there is a demand for developing unbiased, rapid and highly efficient approaches to enrich EVs for diverse applications.

In previous studies, many cell penetrating peptides, such as HIV-1 TAT peptide, antennapedia peptide and other artificially designed peptides, possessed the ability to penetrate the lipid bilayer of the cells, and they have been employed for targeted drug delivery, cancer imaging and other applications^38–40^. Despite possessing the ability to penetrate the cell membrane, few of them have been considered for the isolation of EVs. Among them, a 36-residue polypeptide, namely pH-low insertion peptide (pHLIP), derived from the C helix of bacteriorhodopsin, was discovered in the Engelman’s lab^41^. The wild-type pHLIP (WT pHLIP) is capable of stably inserting into the phospholipid bilayer of cells in an α-helical conformation under an acidic microenvironment^42^. Under neutral conditions, the WT pHLIP assumes nonhelical and peripheral conformations while retaining the ability to bind to the phospholipid bilayer. However, this binding is reversible and unstable under natural biological conditions. That’s the reason that the WT pHLIP is frequently utilized for targeted drug delivery owing to its acid-responsive property^42–44^, but it has been rarely employed for EVs isolation and enrichment^45^. Motivated by the similar structure of EVs with cell membrane, we hypothesized that the pHLIP might be applied for the isolation of EVs, employing the strategy of “old tricks catch new birds”. However, the WT pHLIP was not feasible for isolation of EVs under natural biological conditions. Therefore, it is necessary to reconfigure and modify the WT pHLIP peptide to achieve aforementioned purpose, enabling the peptide transform into a more stable α-helical conformation under neutral conditions and successfully insert into the phospholipid bilayer of EVs. As well-known, enhancing the appropriate hydrophobicity and intramolecular hydrogen bonds of a peptide could be an efficient approach to render it a more stable α-helical conformation^46, 47^, irrespective of the pH effect. Hence, this strategy was adopted to engineer an artificial peptide based on pHLIP for the first time to catering application in rapidly and unbiasedly isolating EVs from complex clinical samples.

We reconfigured the N terminal of Cysteine (Cys) and Alanine (Ala) amino acids instead of Glycines (Gly) from the WT pHLIP and functionalized the **<u>a</u>**rtificial **i**nsertion **<u>p</u>**eptide (AIP) with biotin. The circular dichroism and molecular dynamics simulations revealed that the **<u>b</u>**iotin functionalized **<u>a</u>**rtificial **i**nsertion **<u>p</u>**eptide (BAIP) could form more stable α-helix conformation, feasibly inserting into the membrane of the EVs under neutral conditions. Moreover, the captured EVs could be released and used for subsequent extraction and molecular detection of the cargo in EVs for liquid biopsy (Fig. 1). Compared with the gold standard method (i.e., UC), the BAIP-based system improved the capture efficiency to ∼ 80% with not only reduced isolation time from 4-5 hours to 20 minutes, but also without need for expensive labor or bulky equipment. The isolated EVs were of high purity and could be effectively employed for the proteomics analysis and whole exome sequencing (WES). We successfully used the BAIP coped with the real-time PCR (qPCR) to detect the gene mutations in EVs (e.g., epidermal growth factor receptor (EGFR) and V-Ki-ras2 Kirsten Rat Sarcoma Viral Oncogene Homologue (KRAS) at codons 12 and 13) from the clinical samples. Meanwhile, the experimental results demonstrated that the meticulously-engineered BAIP could also be modified to various micro-nano interfaces for unbiased and efficient capture of EVs overcoming pH limitation. Overall, this novel strategy might be employed to reconfigure and modify many other peptides and proteins for the EVs isolation and extended the diverse applications of EVs, e.g., discovery of new biomarkers on EVs for cancer diagnosis.

**Figure 1.**
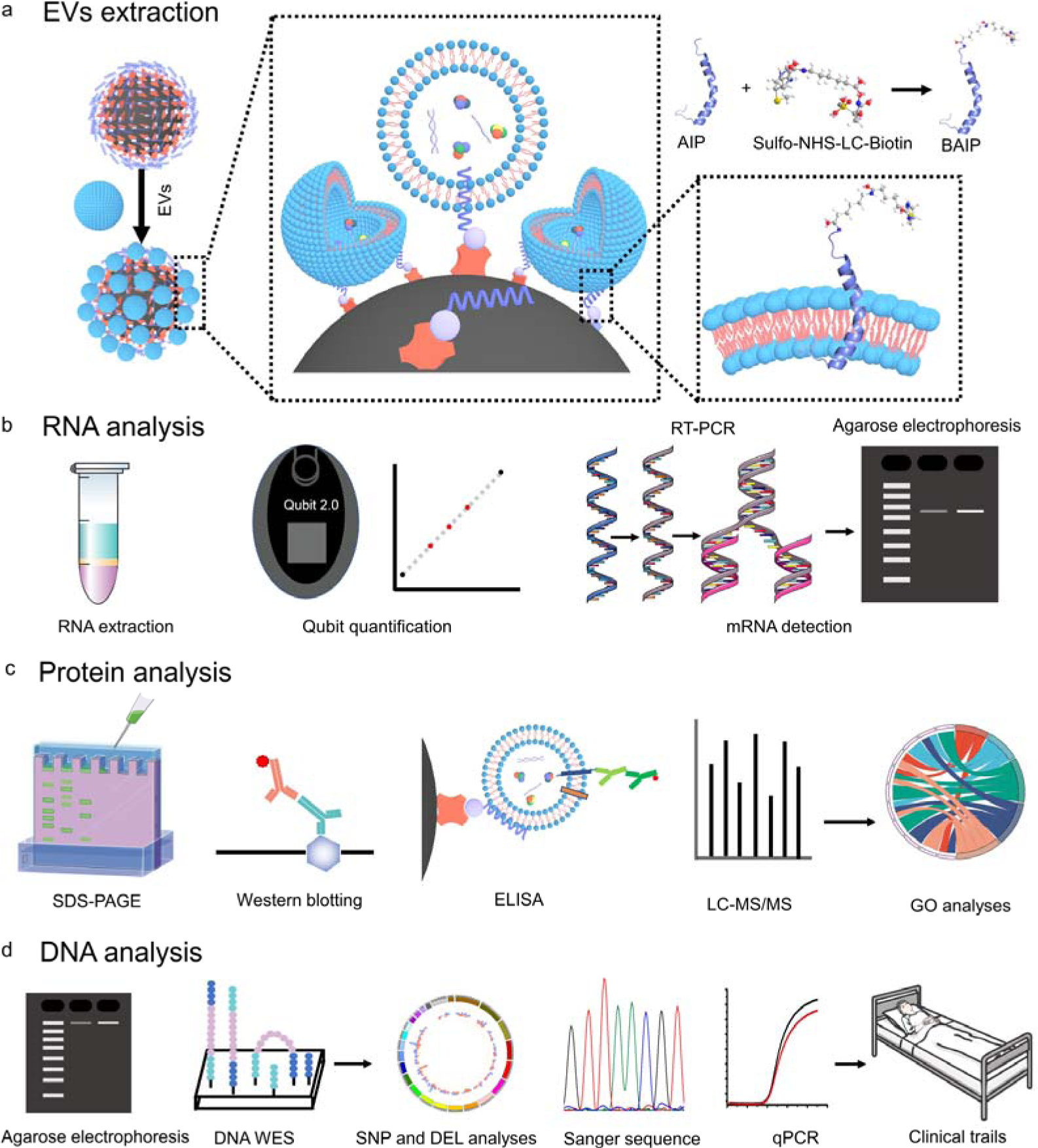
Scheme of the BAIP-based EVs enrichment and downstream analyses. **a**, Scheme of capture of EVs with the BAIP-based method. **b**, Scheme of the extraction of cargo RNA contents from isolated EVs and the subsequent RNA quantification, mRNA detection and RNA agarose electrophoresis characterization. **c**, Scheme of the protein analysis, such SDS-PAGE, Western blotting, enzyme-linked immunosorbent assay (ELISA) and LC-MS/MS. **d**, Scheme of the DNA analysis, such as DNA electrophoresis characterization, WES, sanger sequencing and qPCR for DNA mutation detection.

## Results

### Design, preparation and characterization of the BAIP

In this study, we chose the WT pHLIP as the basic template for reconfiguration and functionalization to isolate EVs. We selected it due to its evolution in nature, forming a transmembrane α-helix at acidic microenvironment (pH<6.5) and spontaneously inserting into the phospholipid bilayer^41, 42^. As mentioned above, we would like to reconfigure and modify the WT pHLIP in order to form α-helical conformation under neutral biological systems. As the appropriate hydrophobicity and intramolecular hydrogen bonds can induce the peptide to form the α-helical conformation, we supposed to change the N-terminal amino acid of the WT pHLIP in order to increase the hydrophobicity and promote the formation of the intramolecular hydrogen bonds of the peptide, thereby facilitating the formation of the α-helical structure under natural pH conditions. Meanwhile, we intended to maintain the stability and unchanged nature of the remaining amino acids. This approach was driven by the fact that the middle segment of pHLIP forms an α-helix conformation, while the C-terminal of pHLIP inserts into the phospholipid bilayer (depicted as the orange and green regions in Supplementary Table 1). Nevertheless, there are still millions of reconfigurations available to enhance the hydrophobicity of the pHLIP. We then calculated the grand average of hydrophilicity (GRAVY) values for the artificial peptide, which reflected their relevant hydrophilic properties. A higher positive GRAVY value indicates a stronger hydrophobic characteristic (Supplementary Table 1). Thus, we meticulously selected a reconfiguration that enhanced the appropriate hydrophobicity of the peptide while maintaining a balance and avoiding excessive hydrophobicity, which could lead to poor water solubility. To achieve this, we replaced the Gly of the WT pHLIP at N-terminal region with Cys and Ala (i.e., AIP, Supplementary Table 2). This transformation changed the R groups of the corresponding amino acids from −H to −CH_3_ and −CH_2_SH, thereby enhancing the hydrophobicity of the peptide. We observed the GRAVY value of the peptide increased from 0.158 (WT pHLIP) to 0.300 (AIP), which might be appropriate for increasing the hydrophobicity of AIP, promoting the formation of α-helix conformation. Moreover, we aimed to modify the N-terminal amino acid of AIP by introducing a biotin moiety (using a Sulfo-NHS-LC-Biotin kit) with a relatively long alkyl chain (Fig. 2a), resulting in a little increase in hydrophobicity. Additionally, the biotin modification could facilitate the formation of intramolecular hydrogen bonds in the AIP, contributing to stabilize the α-helix structure.

**Figure 2.**
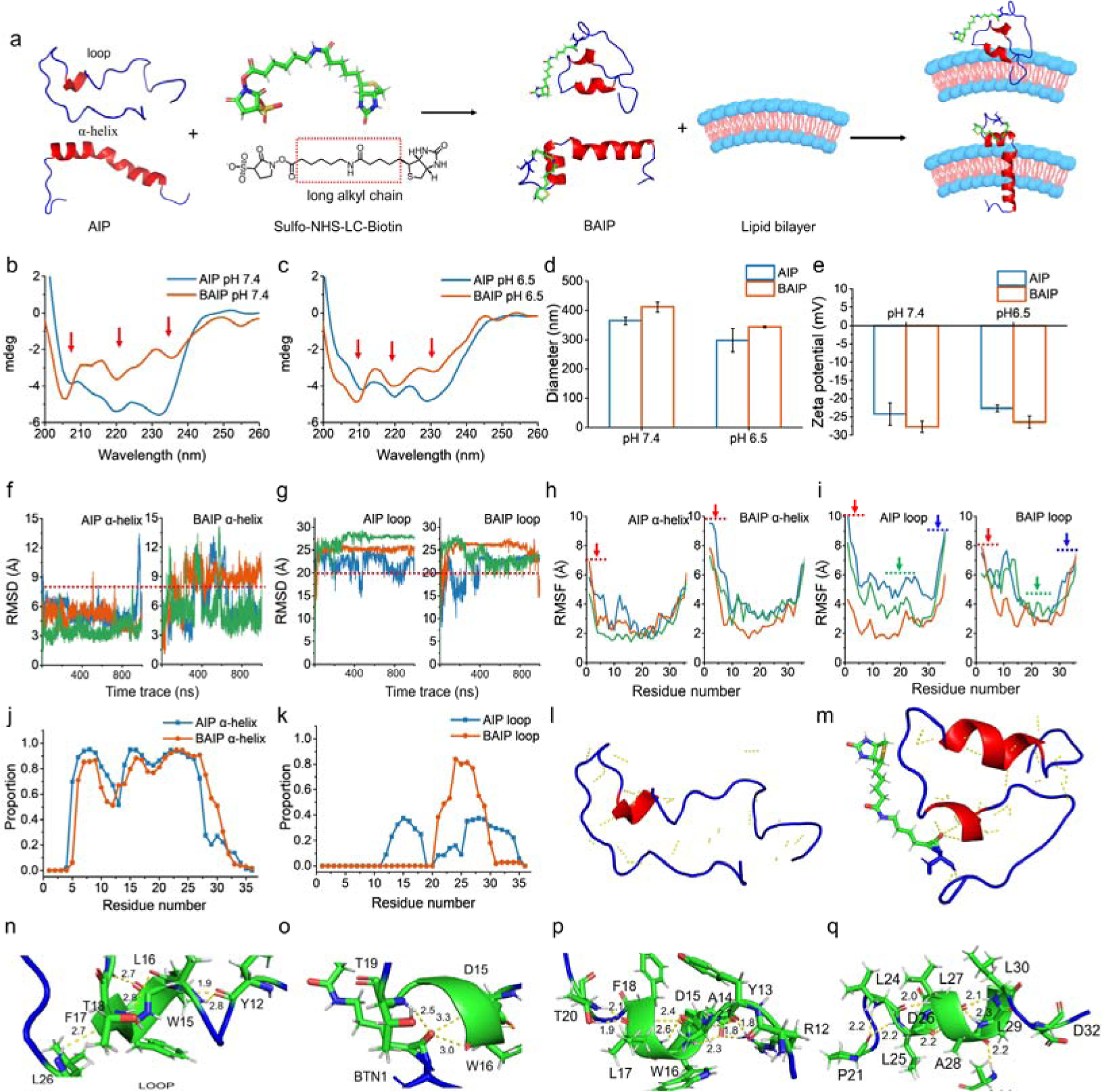
Design and engineering reconfiguration of WT pHLIP to BAIP. **a**, Schematic of AIP modified with biotin and inserted into the phospholipid bilayer. **b**,**c**, Circular dichroism spectra of AIP and BAIP at pH 7.4 and 6.5, respectively. **d**,**e**, Changes in hydration size and zeta potential of AIP and BAIP at pH 7.4 and 6.5, respectively. **f**,**g**, Root Mean Square Deviation (RMSD) of AIP and BAIP with initial helix and loop conformation, respectively. **h**,**i**, Root Mean Square Fluctuation (RMSF) of AIP and BAIP with initial helix and loop conformation, respectively. **j**,**k**, Proportion of each amino acid forming α-helix structure of AIP and BAIP with helix state and loop state. **l**,**m**, Intramolecular hydrogen bonds of reconfigured AIP and BAIP with loop conformation analyzed by PyMOL, respectively. **n**, Intramolecular hydrogen bonds in AIP between residues L16-L18 with loop conformation. **o**, Intramolecular hydrogen bonds in BAIP between Biotin and D15, W16 and T19 with loop conformation. **p**,**q**, The intramolecular hydrogen bonds in BAIP between D15-F18 and L25-L30 with loop conformation, respectively. The unit of hydrogen bond length in Fig. n-q is Å.

In order to verify above hypothesis, we used circular dichroism to measure the conformation of the AIP and BAIP under neutral and weakly acidic conditions (pH 6.5). As depicted in Fig. 2b, the AIP exhibited an α-helix conformation (∼210 nm, ∼220 nm) under neutral condition, which was different from that of WT pHLIP. Additionally, it also showed strong irregular curly structure (∼230 nm). These findings were consistent with the observations under acidic condition (Fig. 2c). After the modification with biotin, the α-helix conformation of AIP was maintained under both conditions. It is noteworthy that the irregular curling structure of BAIP was weakened than AIP (Fig. 2b and 2c), which was consistent with the above hypothesis. Anyway, the almost identical peaks were observed for BAIP under neutral and acidic conditions, implying the BAIP might display similar structure and conformation under both conditions. Meanwhile, the hydrated particle size and zeta potential of the BAIP in acid and neutral phosphate buffered saline (PBS) were increased compared to AIP (Fig. 2d and 2e), which confirmed the successful functionalization of the AIP with biotin.

We then used molecular dynamics simulations to study the changes of the structure and conformation of the AIP and BAIP. According to the results of circular dichroism (Fig. 2b and 2c), the AIP and BAIP exhibited α-helix and loop conformations under both conditions, which were simulated as initial states, and each system was independently simulated with three different trajectories. We observed that the overall structural changes in BAIP were more pronounced compared to AIP, irrespective of whether the initial model was in the α-helix or loop state (Fig. 2f and 2g). This indicated that the biotin modification might interact with AIP (such as hydrogen bonds), resulting in more substantial structural alterations. For more details, when the α-helix state was used as the initial model for simulation, the biotin modification exhibited a more significant impact on the amino acid structure of the N-terminal of AIP. However, it had minimal effect on the middle and C-terminal portions, leading to a significantly stable structure (Fig. 2h and 2i). On the other hand, when the loop state was used as the initial model, the biotin modification had a notable effect not only on the N-terminal amino acid structure, but also on the middle and posterior amino acids of AIP. This effect was observed as reduced flexibility of amino acids, providing the evidence that the biotin modification promoted the formation of secondary structure in AIP (Fig. 2h and 2i).

Furthermore, we utilized the Dictionary of Secondary Structure of Proteins (DSSP) to analyze each amino acid and statistically determine the probability of forming different secondary structures (Supplementary Fig. 1). The statistical results indicated that when the initial state was α-helix state, the AIP had a high probability of forming a helical secondary structure between N5-D31, which remained stable even after the biotin modification (Fig. 2j, Supplementary Table 3 and 4). On the other hand, when AIP was initially in the loop state, apart from amino acids A13-T18 and L26-E34 which had an approximately 30% probability of forming helical secondary structure, the residual amino acids in AIP predominantly exhibited no secondary structure. However, the probability of helical secondary structure formation of BAIP was significantly improved, with segments L24-L27 reaching a helix probability exceeding 80% (Fig. 2k, Supplementary Table 5 and 6).

In addition, the formation of intramolecular hydrogen bonds of AIP and BAIP was analyzed (Supplementary dataset 1). When the initial state of AIP was in a helical conformation, the modification with biotin predominantly resulted in intramolecular hydrogen bonds with N6, Q5, and W10 (Supplementary Fig. 2). In the case of the initial state being a loop conformation, the intramolecular hydrogen bonds were relatively simple, with only three amino acids (L16-L18) forming a helical structure, while the remaining segments lacked any secondary structure (Fig. 2l and 2n, Supplementary Fig. 3a). Following the biotin modification of AIP, the intramolecular hydrogen bonding became more complex. Specifically, amino acids from D15-F18 and L25-L30 formed a helical structure, and biotin formed intramolecular hydrogen bonds with D15, W16, and T19, which contributed to the stability of the helical conformation (Fig. 2m and 2o-2q, Supplementary Fig. 3b). In summary, our AIP exhibited both α-helix and loop conformations under neutral conditions, and the biotin modification facilitates the further formation and stability of the α-helical conformation.

**Figure 3.**
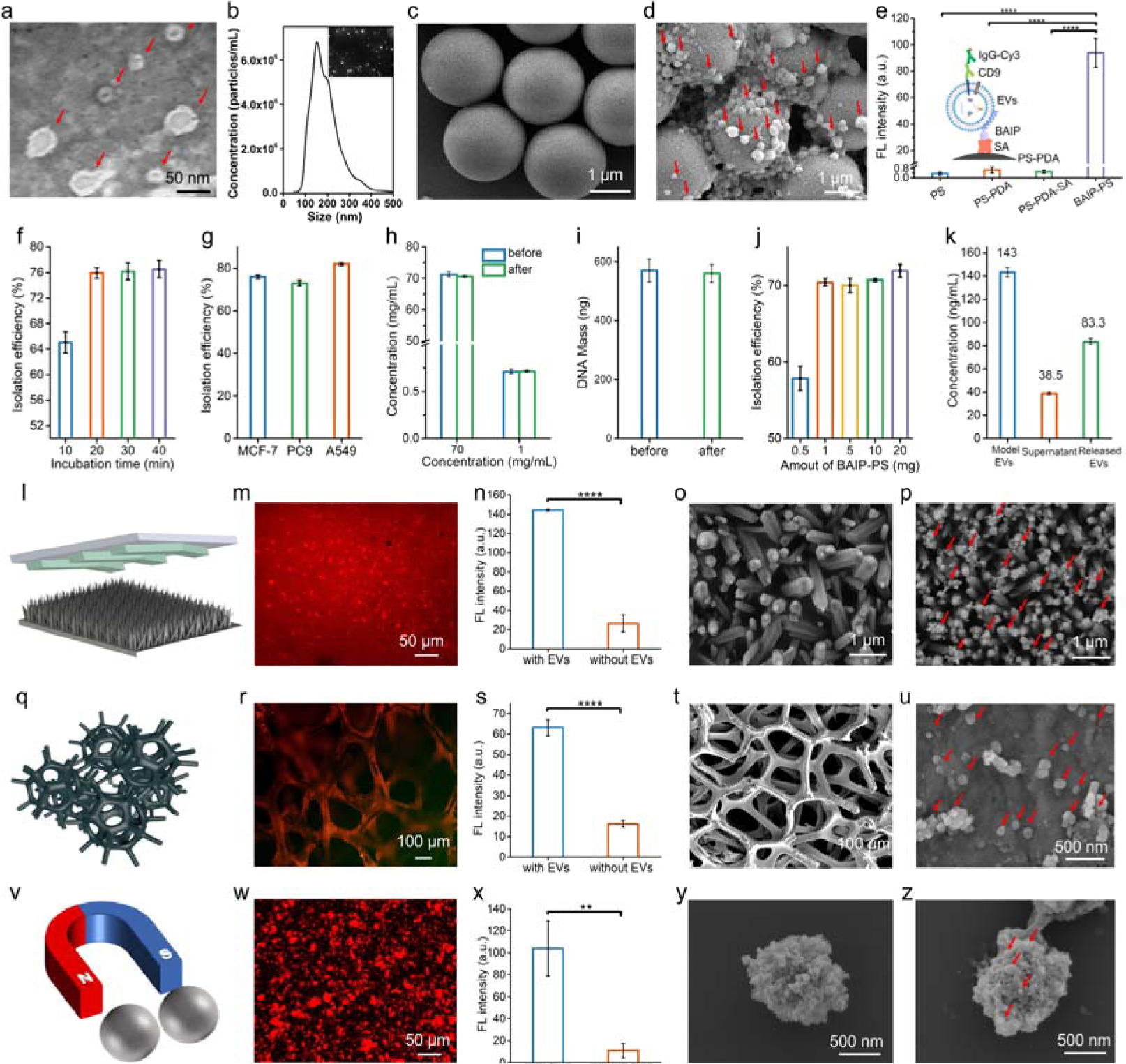
The isolation and release of EVs by BAIP. **a**, TEM image of EVs derived from MCF-7 cells. **b**, NTA curve of EVs derived from MCF-7 cells. **c**, SEM image of BAIP-PS particles. **d**, SEM image of EVs captured on BAIP-PS particles. **e**, Verification of the feasibility of EVs captured on PS, PS-PDA, PS-PDA-SA and BAIP-PS by the immuno-fluorescence experiments. **f**, The influence of incubation time on isolation efficiency of BAIP-PS for model EVs. **g**, The isolation efficiency of BAIP-PS for different types of cells derived EVs. **h**,**i**, Nonspecific adsorption results of BAIP-PS for BSA and DNA, respectively. **j**, The isolation efficiency of EVs from healthy-donor plasma samples as a function of the dosage of BAIP-PS. **k**, The release of the EVs from the BAIP-PS. The concentrations of model EVs, supernatant, and released EVs measured by NTA. **l**,**q**,**v**, The diagram of functionalized interfaces for EVs capture, including ZnO nanorods integrated microfluidic chip (**l**), PDA coated nickel foams (**q**) and SA-functionalized magnetic beads (**v**). **m**,**n**,**r**,**s**,**w**,**x**, The immuno-fluorescence images of EVs captured on different interfaces modified by BAIP and their corresponding fluorescence intensities. ZnO nanorods integrated microfluidic chip (**m**,**n**); PDA coated nickel foam (**r**,**s**); SA-functionalized magnetic beads (**w**,**x**). **o**,**p**,**t**,**u**,**y**,**z**, SEM images of EVs captured on various interfaces functionalized with BAIP (**o**,**t**,**y**) and the corresponding control groups without the addition of EVs (**p**,**u**,**z**). ZnO nanorods integrated microfluidic chip (**o**,**p**); PDA coated nickel foam (**t**,**u**); SA-functionalized magnetic beads (**y**,**z**). Data are represented as mean ± S.D. (n=3). Statistical analysis was assessed by two-tailed Student’s t-tests; \*\**P* < 0.01, \*\*\**P* < 0.001 and \*\*\*\**P* < 0.0001.

### Isolation and enrichment of EVs through BAIP-based system

According to above results, we proceeded experiments to validate the interaction between BAIP and cell membrane. The efficiency of BAIP in inserting into the phospholipid bilayer of cells was evaluated by using the MCF-7 cells as model in PBS buffer at pH 7.4, while Cy3 modified streptavidin (SA-Cy3) was used to characterize the interactions between BAIP and cells. As shown in Supplementary Fig. 4a and 4b, BAIP treated cells displayed a robust fluorescence signal on their membranes as opposed to those untreated cells. Almost all the cells treated with the BAIP exhibited red fluorescence signals, which was further confirmed by flow cytometry analysis (Supplementary Fig. 5). These observations suggested that BAIP could proficiently insert into the phospholipid bilayer of cells under neutral condition, potentially facilitating the separation and enrichment of EVs. Moreover, the binding between BAIP and cells was fast and 20 min incubation was sufficient for the BAIP inserted into the cell membrane (Supplementary Fig. 6a and 6b). We then compared the efficiency of BAIP in inserted into the cell membrane under different pH conditions. The results reveled that the efficiency under neutral condition was comparable to that under acidic condition (Supplementary Fig. 7), supporting the success of our reconfiguration of the WT pHLIP.

Afterward, we tried to employ the BAIP to isolate the EVs in model samples. Firstly, the EVs secreted from MCF-7 cells were isolated through UC, and the obtained EVs were used as a model to evaluate the enrichment performance of EVs using BAIP. The identity of the model EVs was confirmed by transmission electron microscope (TEM). In the TEM image, they displayed a saucer-like morphology, with the diameter ranging from 30 nm to 200 nm (Fig. 3a). Additionally, nanoparticle tracking analysis (NTA) results revealed that the average size of EVs was 193.8 nm, with a concentration of approximately 8.81×10^10^ particles/mL (Fig. 3b).

The BAIP functionalized polystyrene microspheres (PS) were then chosen to unbiasedly separate and enrich EVs from the buffer and complex clinical samples with intricate matrices. The scheme illustrating the modification of BAIP on PS was presented in Supplementary Fig. 8. Initially, the PS were coated with polydopamine (PDA) through self-polymerization of dopamine (DA) under alkaline conditions^48, 49^. This coating served two purposes, facilitating subsequent surface modification and preventing non-specific adsorption. The PS-PDA particles were then subjected to a one-step Michael addition reaction, in which the catechol functional groups on PDA interacted with the amino groups on SA, resulting in the modification of SA onto the PS surface. The optimization of SA modification on the PS was carried out using the immuno-fluorescence method. In this process, biotin-labeled mouse anti-human IgG was first captured on the SA-functionalized PS, followed by the utilization of Cy3-labeled goat anti-mouse IgG for fluorescence imaging to evaluate the degree of SA modification. According to the results in Supplementary Fig. 9, the strongest fluorescence was observed when PS-PDA reacted with 20 μg/mL of SA. which was therefore selected as the optimal condition for PS modification in subsequent experiments. Finally, the monodisperse PS particles functionalized with BAIP (referred to as BAIP-PS) were obtained through the high affinity between SA on PS and biotin in BAIP, with a mean size of 1.9 μm (Fig. 3c). The changes in hydration size and zeta potential observed during the process confirmed the successful modification of BAIP on the PS surface (Supplementary Fig. 10a and 10b). It should be noted that the BAIP-PS exhibited a zeta potential of −38 mV, which might be favorable for reducing non-specific adsorption of negatively charged interfering proteins or other impurities.

We initially validated the feasibility of the BAIP-PS for unbiased isolation and enrichment of the model EVs. The EVs were first captured on BAIP-PS and subsequently imaged by the SEM. The results clearly showed the presence of large amount of EVs (red arrows) on BAIP-PS compared to the BAIP-PS alone (Fig. 3c and 3d), thus supporting the capability of BAIP for isolation and enrichment of EVs. The immuno-fluorescence ELISA experiments were then conducted to prove the feasibility further. CD9, a cluster-of-differentiation protein molecule that is widely and extensively expressed on the EVs, was selected as a target protein to verify the isolation of EVs by BAIP. As shown in Supplementary Fig. 11, the captured EVs were further recognized by anti-human CD9 antibody and identified by Cy3-labeled anti-mouse IgG. The results indicated that the strong fluorescence was exclusively detected on the BAIP-PS, whereas there was no notable fluorescence observed in the material groups without the addition of EVs (Supplementary Fig. 12). Notably, the fluorescence intensity on the BAIP-PS was significantly different from that of the control groups, implying the crucial function of BAIP in the capture of EVs (Fig. 3e).

Next, the capture parameter of BAIP-PS towards the EVs was optimized and the capture efficiency was evaluated. It should be noted that no standard method is available for EVs quantification so far. The current methods for EVs detection, such as NTA and total protein detection, are limited by their failure to specifically identify EVs and interference from non-exosomal particles or proteins. As an alternative, the detection of RNA in EVs is considered to be a more reliable method, as RNA molecules are protected by lipid membranes, leading to enhanced stability. In contrast, the cell-free RNA molecules in plasma are susceptible to degradation by various RNA-degrading enzymes in the bloodstream, but potentially not interfering with the detection of RNA in EVs. Taking the advantages of stable RNA in EVs, the detection of RNA was adopted for quantification of EVs in our experiments. The isolation efficiency was determined by calculating the mass fraction of RNA extracted from the captured EVs over that obtained from the total EVs. As depicted in Fig. 3f, the isolation efficiency increased from 65.1% to 76.5% with the incubation time increasing from 10 min to 1 h for capture of EVs by BAIP-PS and reached a plateau at 20 min. This observation was also consistent with the results obtained from the interaction of BAIP with the cells (Supplementary Fig. 6), further demonstrating the fast interaction of BAIP with phospholipid bilayer even in this fixation status. Hence, 20 min was chosen for subsequent experiments.

To confirm the versatile and unbiased enrichment of EVs by BAIP, we utilized the peptide to capture EVs derived from various cell types. Specifically, we selected EVs derived from MCF-7, PC9, and A549 cells, which exhibited high, medium, and low EpCAM expression levels respectively, to assess the capture efficiency of EVs by BAIP. The results revealed that the isolation efficiencies of EVs from MCF-7, PC9 and A549 cells were 76%, 72% and 82%, respectively (Fig. 3g). These findings suggested that our proposed method appeared to be capable of capturing EVs from various cell subpopulations with less bias, leading to an improved enrichment in EVs.

We further investigated the anti-inference capability of the BAIP towards the protein and nucleic acid as actual plasma/serum samples contain diverse proteins and short DNA fragments that may potentially impact the capture of EVs. As shown in Fig. 3h and 3i, despite whether the presence of high (70 mg/mL)/ low (1 mg/mL) level of proteins or DNA fragments (∼160 bp, 570 ng), the capture efficiency of them by the BAIP-PS was negligibly (<1%). These findings supported that there was no discernible interaction between the BAIP and proteins or DNA. The negatively charged BAIP-PS inhibited the adsorption of negatively charged proteins and DNA as a result of the electrostatic repulsion. We hypothesized that the similar phenomenon may extend to other cellular components, such as globulins and fibrinogens with negative charges at physiological pH. Additionally, it was worth noting that the PDA coating as well as BAS blocking might also synergistically inhibit non-specific adsorption of albumin, cell-free DNA, and other contaminants^49^.

With excellent performance towards model EVs, the BAIP-PS was then attempted to isolate EVs from plasma samples. In our previous study, we found that the albumin could interfere with the insertion of lipid-nanoprobe (LNP, DSPE-PEG), into the membranes of EVs, leading us to increase the quantity of LNP for the isolation of EVs in 100 μL of plasma^36^. However, even with excessive lipid probes (200 nmol), we only achieved a 48.3% isolation efficiency of EVs in plasma. Doubling the amount of the LNP only slightly increased isolation efficiency to 49.5% (not statistically significant; *P* > 0.05, two-tailed t-test). In contrast, we found that using 1 mg of our BAIP-PS to enrich EVs in 100 μL of plasma resulted in an isolation efficiency of 70%, which was slightly lower than that achieved by isolating EVs in PBS buffer (76%). Interestingly, the isolation efficiency of EVs in plasma remained constant at 70-72% regardless of the dosage of BAIP-PS from 1 mg to 20 mg (Fig. 3j). This could be due to the limitations in reaction dynamics equilibrium between nano-sized EVs and micro-sized PS particles^50, 51^. Nevertheless, our proposed BAIP-PS based method achieved superior stability and reliability in isolation efficiency compared to the LNP-based unbiased EVs isolation method, making it potentially suitable for the clinical requirements.

The BAIP was also utilized to modify a variety of interfaces for the efficient and unbiased capture of EVs. These interfaces included ZnO nanorods integrated microfluidic chips, PDA coated nickel foams, and SA-functionalized magnetic beads (Fig.3l, 3q and 3v, Supplementary Fig. 13 and 14). The successful capture of EVs was initially confirmed through immuno-fluorescence assays, where the CD9 biomarker on the captured EVs was measured (Fig. 3m, 3r, and 3w). The fluorescence signals of the BAIP-functionalized interfaces were significantly distinguishable from those of the control groups without BAIP modification (Fig. 3n, 3s, and 3x). Meanwhile, SEM results clearly revealed the captured EVs on these BAIP-functionalized interfaces (Fig. 3p, 3u, and 3z) in comparison to the interfaces themselves (Fig. 3o, 3t, 3y). Our findings indicated that the diverse BAIP-functionalized interfaces could efficiently and unbiasedly isolate and enrich EVs.

In light of the multiple applications of EVs, including drug delivery and wound healing, release of the captured EVs was equally necessary. To achieve this, we employed Sulfo-NHS-SS-Biotin, a molecule containing a disulfide bond, to modify AIP (referred to as ss-BAIP, Supplementary Fig. 15). This modification facilitated the release of the captured EVs by cleaving the disulfide bond between Biotin and AIP using dithiothreitol (DTT). Immuno-fluorescence analysis showed that fluorescence signal on BAIP-PS disappeared after the release of fluorescence labelled EVs (Supplementary Fig. 16). The release efficiency was calculated to be 79.5% ±2.6% (Fig. 3k), indicating that the captured EVs could be satisfactorily released and utilized in subsequently functional assays.

### Detection of proteins and nucleic acids in EVs samples

The BAIP-functionalized interfaces facilitate the isolation and enrichment of EVs on their surfaces, allowing for subsequent molecular analyses with the profiling of proteins and nucleic acids. We first validated that the EVs isolated by BAIP could be used for protein analysis. Initially, the proteins on the captured EVs through BAIP-PS and UC techniques were examined and compared through the SDS-PAGE. The outcome revealed numerous diffuse bands, ranging from 25kD to 180kD, in the EVs obtained from both methodologies (Supplementary Fig. 17a). It is noteworthy that no significant difference was observed when comparing the protein bands of the isolated EVs through the BAIP-PS based approach (Lane 3) with those of the model EVs obtained by the conventional UC method (Lane 2). The Western blot technique was next used to identify two specific membrane proteins, CD9 and glyceraldehyde 3-phosphate dehydrogenase (GAPDH, housekeeping protein), on the model EVs and isolated EVs. The findings revealed comparable CD9 and GAPDH bands, as photographed in Fig. 4a and Supplementary Fig. 17b, indicating that our proposed isolation method would not compromise the subsequent protein analysis of EVs.

**Figure 4.**
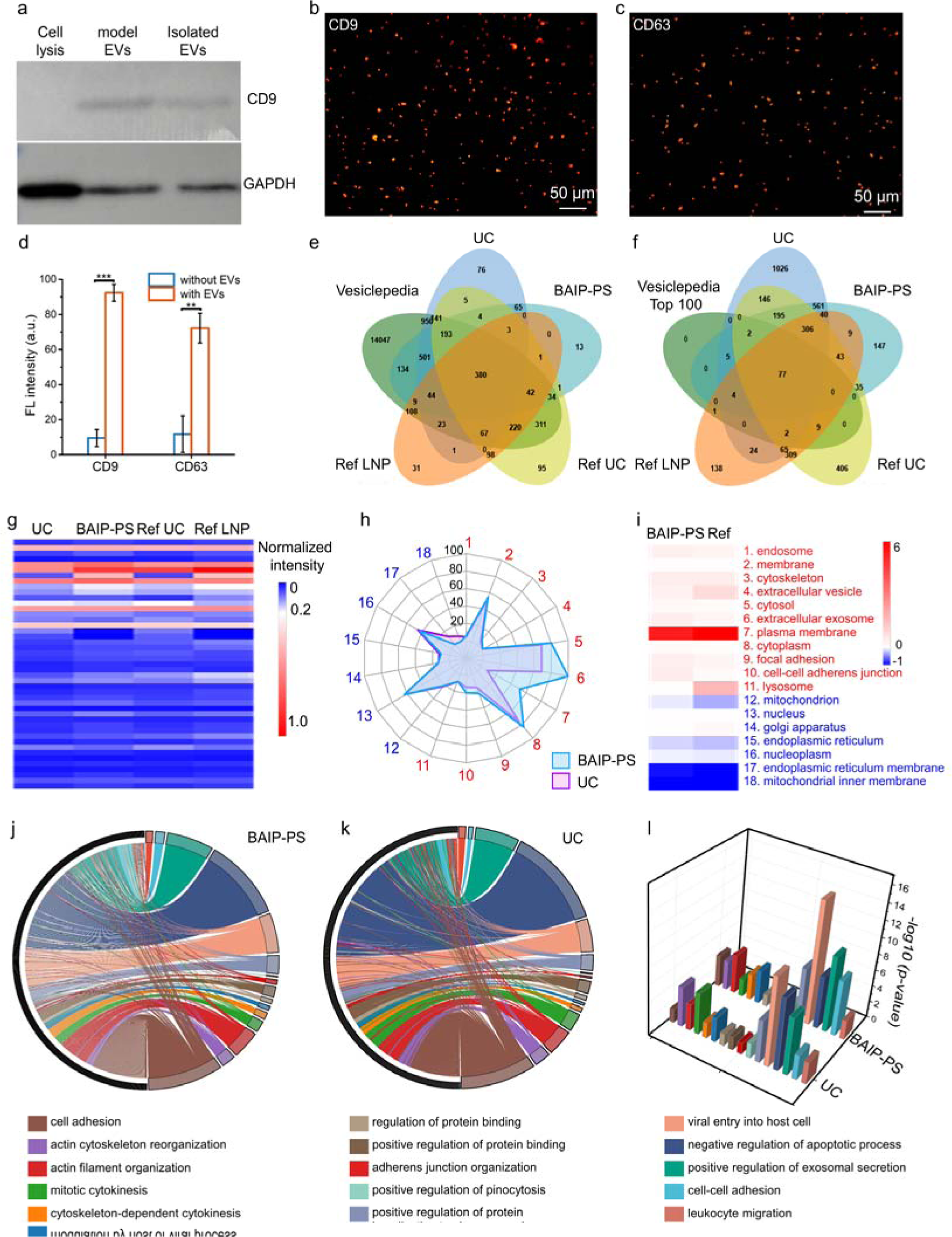
Protein analyses of EVs isolated by BAIP-PS. **a**, Western blotting identified CD9 and GAPDH proteins on the cells and EVs. These proteins were extracted from MCF-7 cells, model EVs and isolated EVs by BAIP-PS. **b**-**d**, Detection of CD9 and CD63 proteins on the isolated EVs by our BAIP-based method. The captured EVs on the BAIP-PS were recognized with either anti-human CD9 antibody, or biotin-CD63 aptamer, which specifically reacted with CD9 or CD63 proteins on EVs. Subsequently, the Cy3-labeled anti-mouse IgG or Cy3-labeled SA was used to bind with CD9 antibody or CD63 aptamer, generating a fluorescent signal for further analysis. The corresponding fluorescence intensity of the CD9 and CD63 proteins on the isolated EVs was calculated by Image J software (**d**). **e**,**f**, Venn diagram showing the overlap among our MCF-7 EVs proteomic data from UC, BAIP-PS, proteomic data of nEV derived from human MDA-MB-231 cells in our previously published report^36^, and public EV database Vesiclepedia (**e**) or Top 100 proteins in the public EV database Vesiclepedia (**f**). **g**, Protein proportion of GO cellular components in proteomics. **h**, Radar chart showing the protein proportion of GO cellular components involved in the composition proteins and the interference proteins of EVs from BAIP and UC-based method. **i**, Proportion of protein changes of GO cellular components involved in the composition protein and the contaminating protein of our method and Ref. [36]. **j**,**k**, Chord of GO analyses for biological processes involved in tumor progression obtained by EVs from BAIP-PS and UC-based method, respectively. **l**, Correlations between the proteome of EVs and tumor regulation pathways from BAIP-PS and UC-based methods. GO analyses for biological process were performed by DAVID database (https://david.ncifcrf.gov/home.jsp). *P* value for the protein enrichment in annotated GO terms (compared to the human genome background) was determined by the EASE Score. Data are represented as mean ± S.D. (n = 3). Statistical analysis was assessed by two-tailed Student’s t-tests; \*\**P* < 0.01, \*\*\**P* < 0.001.

Afterward, the immuno-fluorescence method (Supplementary Fig. 11) was used to detect the proteins on isolated EVs, taking the tetraspanin proteins, CD9 and CD63, as representative proteins. In the presence of EVs, Fig. 4b and 4c exhibited robust fluorescence signals on the BAIP-PS particles, whereas the control groups without EVs demonstrated almost no fluorescence signals (Supplementary Fig. 18). The fluorescence intensities of CD9 and CD63 on the BAIP-PS were significantly different from those of the control groups without EVs (Fig. 4d), indicating that the captured EVs enabled the excellent detection of constituent proteins on BAIP-PS.

Encouraged by the good performance of BAIP-PSto isolate EVs, we used the liquid chromatography-tandem mass spectrometry (LC-MS/MS) to identify potential exosomal proteins, with the comparison among that from UC (Supplementary, dataset 2), our LNP-based method ^36^ and the EVs proteins listed in the publicly available database Vesiclepedia (www.microvesicles.org). The Venn diagram illustrated the overlap between proteomic data of EVs from BAIP-PS, UC and LNP-based methods, as well as Vesiclepedia (Fig. 4e and 4f). Approximately 93.9% of the proteins in EVs isolated by our BAIP-PS based method was identified in 17,204 EV cargo proteins listed Vesiclepedia, while 93.7% of the proteins in EVs isolated by UC was identified. However, in our previously reported LNP-based method, around 87.0% proteins in EVs were identified in the database. When considering the top 100 proteins in the database, our proposed method was able to identify 93 proteins in EVs, while UC and LNP-based methods found 90 and 88 proteins, respectively, indicating a slightly higher efficiency in identifying cargo proteins in EVs by using our proposed method. Our LC-MS/MS data were also compared with the LNP-based methods on 30 key proteins in EVs (Supplementary Table 7) and the results revealed that similar exosomal proteins were found in both methods.

The gene ontology (GO) analysis for cellular components of the identified proteins of EVs from BAIP-PS was further conducted and compared with those from UC-based method and the reported LNP-based method. As depicted in Fig. 4g, the exosomal proteins obtained by BAIP-PS were comparable to those obtained by other methods. Furthermore, the changes of cellular components of the identified proteins from BAIP-PS and UC showed high consistency with those from LNP-based method in our previous report^36^. Meanwhile, recent studies have demonstrated that the protein subpopulations in EVs mainly originated from the cytoplasm, cell membrane and extracellular domain, rather than from subcellular organelles or cell nucleus^52^. These findings provided a theoretical foundation for assessing the purity of EVs based on proteomics. Hence, the proteomes of EVs from both BAIP-PS and UC-based methods were further systematically analyzed and the levels of purity were evaluated and compared. The Radar chart showed the protein proportion of GO cellular components present in both the composition protein and the interference proteins of EVs obtained from the BAIP-PS and UC-based methods (Fig. 4h and Supplementary Table 8). In the chart, the red numbers 1 to 11 represent the composition proteins of EVs, whereas the blue numbers 12 to 18 represent the interference proteins. As shown in Fig. 4h, the larger areas in the red region and smaller areas in the blue region demonstrated that our BAIP-PS based method identified more composition proteins of EVs and fewer interference proteins compared to the UC. This result was consistent with our previous LNP-based method (Supplementary Fig. 19), which also showed similar changes (verse UC-based method) in protein proportions of GO cellular components (Fig. 4i), as evidenced by the red and blue markers indicating composition proteins and contaminant proteins, respectively (Supplementary Table 8). These findings suggested that our BAIP-PS-based method could isolate EVs of higher purity than UC, which may facilitate downstream analysis of EVs accurately and reliably, including tumor screening or diagnosis based on proteomics of EVs.

Moreover, EVs may serve as an indicator for predicting the tumor progression. For this purpose, the GO biological process analyses for proteins of EVs obtained through BAIP-PS and UC-based methods were conducted. Sixteen crucial biological process terms associated with tumor progression were then particularly examined and the corresponding chords from EVs isolated by BAIP-PS and UC-based methods were illustrated in Fig. 4j and Fig. 4k, respectively. After comparing the results, we found that our proposed BAIP-PS based method could recognize 27% more of special tumor-relevant progression proteins in contrast to that of the UC-based method. Meanwhile, the correlations between the proteome of EVs and tumor regulation pathways were determined by the EASE Score (*P value*). Notably, higher correlations in tumor regulation pathways were achieved in our BAIP-PS based method than the UC-based method, with 10 out of 16 crucial biological process terms associated with tumor progression showing either comparable or significant improvement (Fig. 4l). These findings suggested that our BAIP-PS based method for isolating EVs resulted in a significant enrichment of tumor-related proteins that were actively involved in tumor development and progression.

The obtained EVs by BAIP could be used not only for protein analyses, but also for nucleic acid analyses. On the one hand, the ability to detect mRNA in EVs was examined using EGFR and CK19 transcripts as examples. The RNA in isolated EVs was extracted and then the target mRNAs (EGFR and CK19) were amplified with the reverse transcription PCR. As Fig. 5a depicted, the bands of 285 bp fragments of the EGFR coding region and 346 bp fragments of the CK19 coding region were clearly observed on the 2% agarose gel which indicated that the EVs isolated by our BAIP based method could be used for subsequent mRNA analyses. On the other hand, DNA was extracted from the isolated EVs with our BAIP based method followed by 2% agarose gel electrophoresis. Fig. 5b showed the presence of long fragments of DNA in extracted DNA (>10 kbp), which differed from circulating cell-free DNA with a typical apoptotic DNA ladder^53^. Afterward, DNA from the isolated EVs with BAIP-PS and model EVs (PC9 cells derived) were analyzed by WES. DNA from EVs isolated by the two methods spanned all chromosomes (Fig. 5c). The DNA content of EVs isolated by BAIP-PS resembled that of EVs isolated by UC, as indicated from the copy number variation (CNV) plots of the EVs sample from both methods (Fig. 5c). The DNA contents of EVs isolated by UC and BAIP-PS were similar, with a Pearson correlation coefficient of 0.92. The SNP and Del density distribution were analyzed. The density map of SNP revealed a non-random distribution. EVs isolated using UC exhibited higher SNP density in chr3, chr11 and chr19, while EVs isolated with BAIP-PS displayed elevated SNP density in chr1, chr3 and chr11 (Fig. 5d and 5e). At the same time, the mutations of DNA from EVs isolated by UC and BAIP-PS were annotated and summarized. The results showed that the mutations were divided into six types according to their effects on protein coding. Frame_Shift_Del was the most common type in both UC and BAIP-PS based methods (Fig. 5f). Moreover, there were more DEL mutations than SNP mutations in both EVs isolated by UC and BAIP-PS (Fig. 5g). The SNV mutations were divided into six types, among them, T>C and C>T were the most common SNV mutation types in both EVs isolated by UC and BAIP-PS (Fig. 5h).

**Figure 5.**
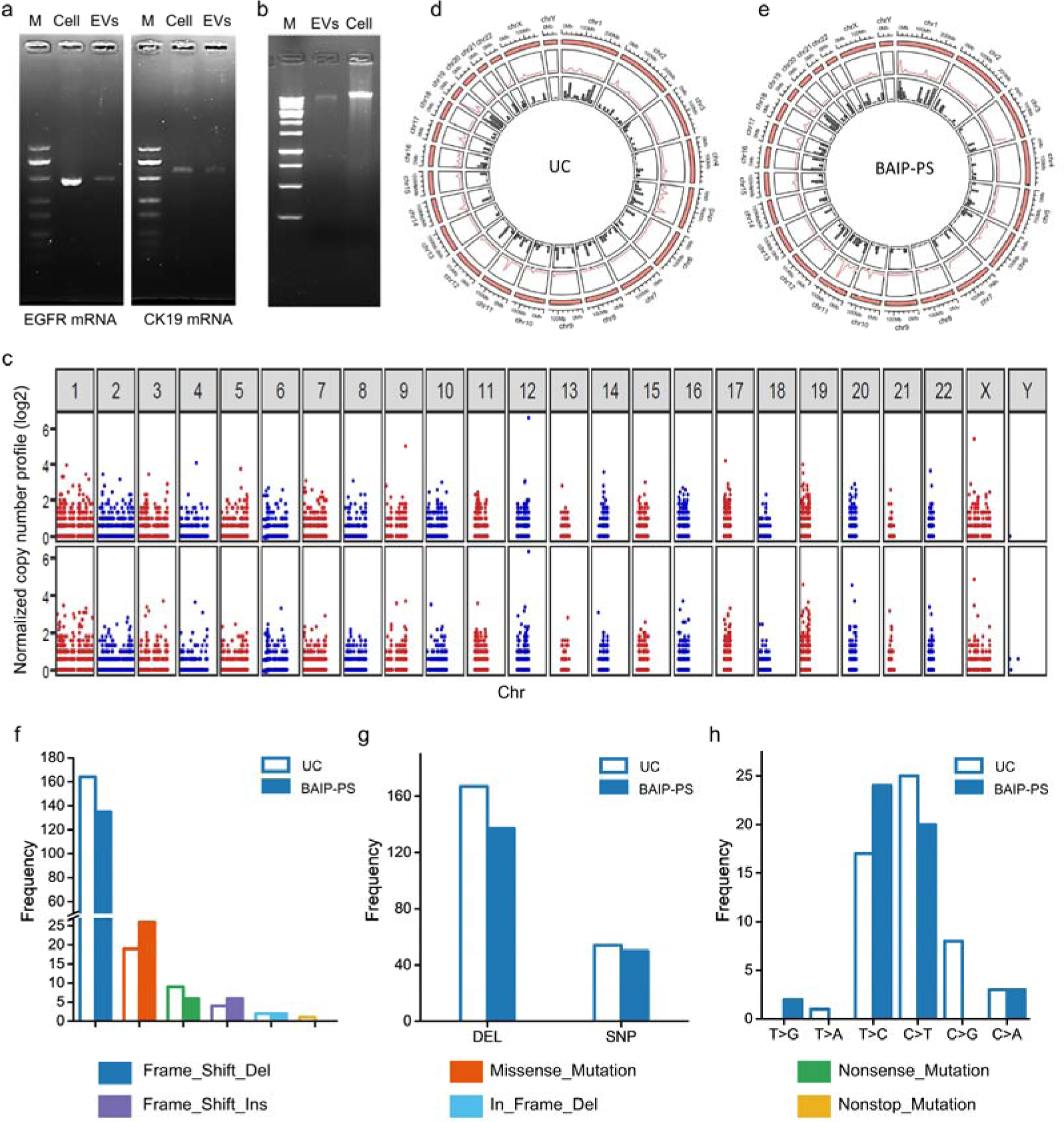
The nucleic acid analyses of EVs isolated by BAIP-PS. **a**, Agarose gel electrophoresis of the fragments from reverse transcription PCR amplification of EGFR mRNA (left) and CK19 mRNA (right). (M, 500 bp DNA ladder. The sequences of the fragments of EGFR and CK19 coding region were listed in Supplementary Table 9. **b**, Identification of DNA extracted from isolated EVs and MCF-7 cells by 2% agarose gel electrophoresis. DNA ladder: 1 kbp. **c**, Plot of the exon-wide copy-number profiles of PC9 EVs isolated with UC (top) and PC9 EVs isolated with BAIP-PS (bottom) generated using Control-FREEC. **d**,**e**, Circos plots of DNA from PC9 EVs isolated by UC and BAIP-PS, respectively. The outer ring is chromosome length, the second layer is SNP density line chart, and the innermost layer is Indel density histogram. **f**-**h**, Bar charts of frequency distribution with different variant classification, different variant types and different SNV mutation types from UC and BAIP-PS based isolation methods, respectively.

### Detection of mutated DNA in EVs from the plasma samples

In order to promote BAIP-based method for EVs isolating to clinical trials, molecular biology methods were used to detect gene mutations in EVs directly. Firstly, 1 μL of the model EVs derived from PC9 cells was added into 99 μL of the plasma from the healthy donors to simulate clinical samples, which were used to test the performance of the gene mutation detection. qPCR assay was implemented to detect the EGFR deletion mutation in exon 19 (EGFR^19Del^). As shown in Fig. 6a, the EGFR^19Del^ mutation in the extracted DNA from isolated EVs by our method was successfully identified with qPCR. Secondly, EGFR-mutant-enriched nested PCR was implemented to enhance the sensitivity for Sanger sequence analysis of EGFR^19Del^ mutation. The desired PCR products of EGFR^19Del^ mutation were detected on the agarose gel (Fig. 6b) and the EGFR^19Del^ mutation was successfully identified by the Sanger sequencing analysis (Fig. 6c). These findings suggested that our BAIP-based method for EVs isolation had the potential to be employed for DNA mutation detection in clinical samples.

**Figure 6.**
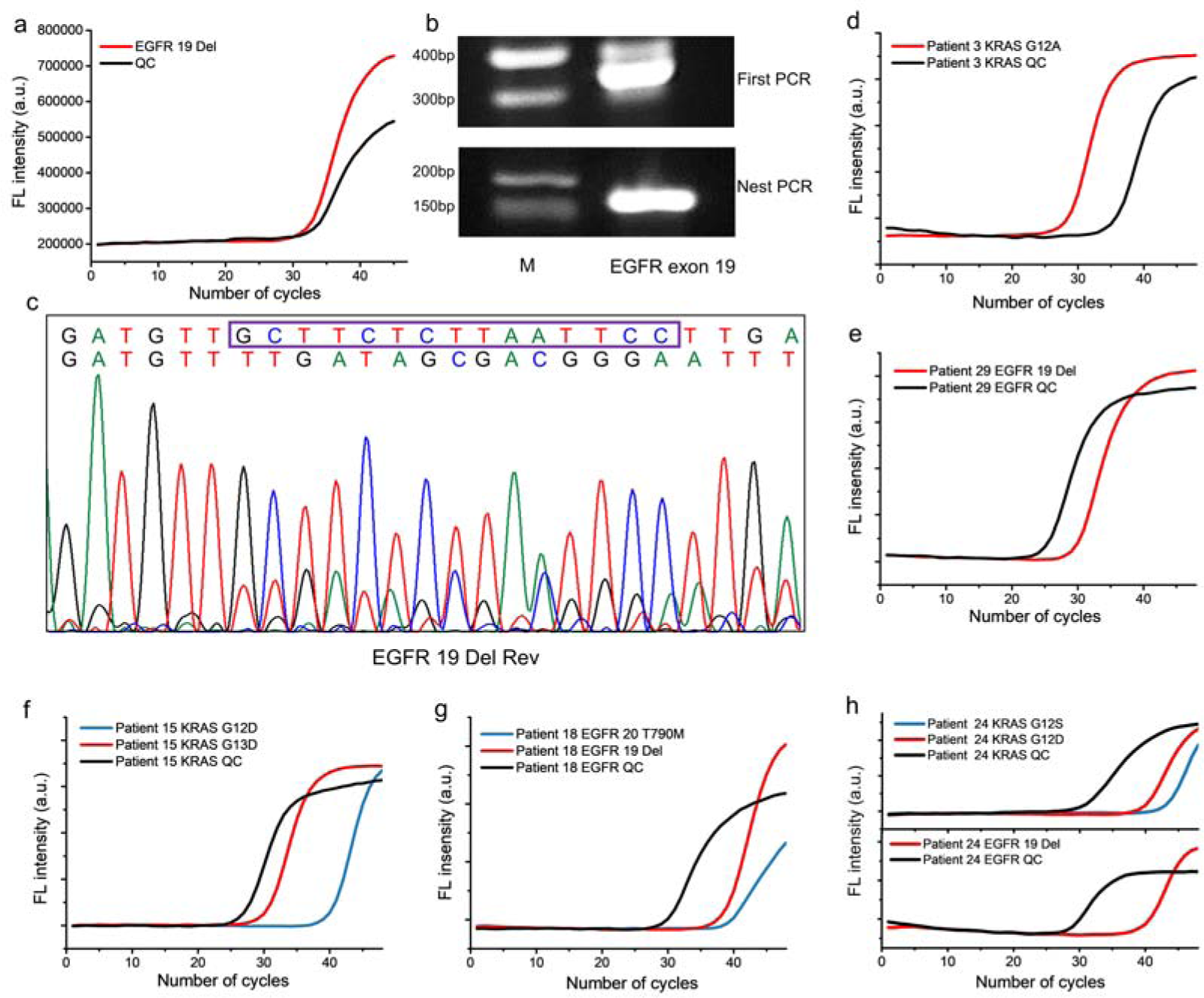
Detection of exosomal DNA mutations from isolated EVs with various methods. **a**, Detection of EGFR^19Del^ mutation from isolated EVs by BAIP-PS in a simulated clinical sample by qPCR. **b**, Gel electrophoresis analysis of EGFR DNA fragments from isolated EVs after a first PCR amplification (top) and a second EGFR-mutant-enriched nested PCR (bottom). M, 500-bp DNA ladder. **c**, Identification of EGFR^19Del^ mutation in isolated EVs from a simulated clinical sample by Sanger sequence. **d**-**h**, qPCR profiles for KRAS mutations and/or EGFR mutations in the isolated EVs by BAIP-PS from patients 3, 29, 15, 18 and 24, respectively.

Each aliquot of 500 μL genuine plasma samples collected from 31 patients and 14 healthy donors was directly processed with BAIP-PS and the DNA was extracted from the isolated EVs to examine the EGFR and KRAS mutations via qPCR. Meanwhile, the mutation detection results of qPCR in patients’ samples were compared with those obtained from next generation sequencing (NGS) or qPCR measurements in the corresponding tissue samples. The results are listed in Table 1 and Supplementary Table 10. The mutation analyses revealed the presence of KRAS G12A, KRAS G12V, KRAS G12D and KRAS G13D point mutations, EGFR^19Del^ mutations, EGFR^18G719C^ mutations in patients 3, 4, 7, 9, 15, 18, 24, 25 and 29, as well as WT KRAS in patients 17, 18, 19, 21, 26, 27 and WT EGFR in patients 2, 30 and 31, which matched well with the results of corresponding tissue samples of patients (Fig.6d-6h, Supplementary Fig. 20). It was noteworthy that these mutations have been previously linked to the development and progression of various types of cancer, making them potentially valuable for guiding accurate personalized medical treatment. Moreover, we identified EGFR^19Del^ mutations in patients 6, 11, 12, 17 and 20 (Supplementary Fig. 21). However, these mutations were not verified through NGS or qPCR analysis of their corresponding tissue samples, due to the limited quantity of samples. On the other hand, despite of our efforts, we failed to detect some mutations in patients, such as 1, 8, 16, 19 and 28, while the NGS or qPCR analysis of tissue identified them, presumably due to the low abundance of cancer-relevant EVs in the plasma samples or due to the alterations in the mutation status between the original tissue biopsy and blood collection. Notably, the EGFR^19Del^ mutations in patient 9, 22, 25 and 27 and EGFR^20T790M^ mutations in patient 18 and 27 were detected. Moreover, KRAS G12D point mutations were detected in patients 15and 24, and KRAS G12S point mutations were found in patients 22 and 24 (Supplementary Fig. 22). However, no EGFR or KRAS mutations were detected in the tissue through NGS or qPCR analysis. This finding demonstrated that our method could detect undetectable mutations with low concentration in tissue. In addition, all KRAS and EGFR alleles in healthy donors were detected as wild type following EVs isolation via the BAIP-PS based method. Nonetheless, the identification of EGFR and KRAS mutations in some patients underscores the potential utility of our assay for mutation analyses in clinical settings.

**Table 1.** KRAS and EGFR mutations in samples from patients.

| Patient number | Sex | Age | Tissue source | KRAS mutation |  | EGFR mutation |  |
| --- | --- | --- | --- | --- | --- | --- | --- |
|  |  |  |  | EVs | Tissue | EVs | Tissue |
| 1 | M | 63 | Lung | NA | WT | WT | exon 21 L858R |
| 2 | M | 67 | Lung | WT | G12R | **WT** | **WT** |
| 3 | F | 76 | Colon | **G12A** | **G12A** | NA | NA |
| 4 | M | 52 | Colon | **G12V** | **G12V** | NA | NA |
| 5 | M | 74 | Lung | NA | WT | WT | exon 21 L858R |
| 6 | F | 67 | Liver | NA | NA | exon 19 Del, exon 20 Ins | NA |
| 7 | M | 75 | liver | **G12D** | **G12D** | NA | NA |
| 8 | F | 68 | Rectum | WT | G12D | NA | NA |
| 9 | M | 56 | Lung | NA | WT | **exon 18 G719X**, exon 19 Del | **exon 18 G719C** |
| 10 | M | 77 | Lung | NA | WT | WT | exon 21 L858R |
| 11 | M | 44 | Liver | NA | NA | exon 19 Del | NA |
| 12 | F | 59 | Liver | NA | NA | exon 19 Del | NA |
| 13 | F | 82 | Lung | NA | WT | WT | exon 21 L858R |
| 14 | M | 48 | Lung | NA | WT | WT | exon 21 L858R |
| 15 | M | 59 | Colon | G12D, **G13D** | **G13D** | NA | NA |
| 16 | F | 83 | Lung | NA | WT | WT | exon 21 L858R |
| 17 | M | 73 | Colon | **WT** | **WT** | exon 19 Del | NA |
| 18 | M | 57 | Lung | **WT** | **WT** | **exon 19 Del**, exon 20 T790M | **exon 19 Del** |
| 19 | M | 76 | Lung | **WT** | **WT** | WT | T790M |
| 20 | M | 63 | Colon | WT | NA | exon 19 Del | NA |
| 21 | F | 62 | Rectum | **WT** | **WT** | NA | NA |
| 22 | M | 45 | Lung | G12S | WT | exon 19 Del | WT |
| 23 | M | 79 | Colon | WT | G12V | NA | NA |
| 24 | M | 50 | Lung | G12S, G12D | WT | **exon 19 Del** | **exon 19 Del** |
| 25 | M | 66 | Colon | **G12A** | **G12A** | exon 19 Del | WT |
| 26 | M | 61 | Liver | **WT** | **WT** | NA | NA |
| 27 | F | 59 | Lung | **WT** | **WT** | exon 19 Del, T790M | WT |
| 28 | F | 59 | Lung | NA | WT | WT | exon 21 L858R |
| 29 | M | 73 | Lung | NA | NA | **exon 19 Del** | **exon 19 Del** |
| 30 | M | 49 | Lung | NA | NA | **WT** | **WT** |
| 31 | M | 53 | Lung | NA | WT | **WT** | **WT** |

Bold text indicated that the mutation of EVs was consistent with that of tissue. NA, not available. WT, wild type. Del, deletion.

## Discussion

The efficient separation and enrichment of EVs from body fluids is crucial for various biomedical applications, including molecular detection for cancer diagnosis, functional application in regenerative medicine, and drug carriers for cancer therapy^22, 54, 55^. However, currently, there are few of rapid, efficient and low-cost methods to isolate and enrich EVs with high purify^1, 17, 56^. Meanwhile, the EVs display heterogenous properties in terms of their membrane proteins and nucleic acids^35^. Thus, conventional approaches for EVs isolation and detection, such as those relying on specific antibodies or aptamers-based techniques, frequently encounter biases and fall short in capturing disease-relevant EVs. In addition, for the cancer biomarker screening, the biased approaches for EVs isolation and enrichment may potentially lead to the loss of cancer-related information, despite that the subsequent unbiased omics analyses, such as proteomics and transcriptomics, are performed.

Thus, in this study, we provided a novel strategy to reconfigure and modify the membrane-penetrating peptide to achieve rapid, cost-effective and unbiased isolation and enrichment of EVs by inserting the modified peptide to phospholipid bilayer of EVs. We changed the amino acid residue at the N-terminal of pH-sensitive WT pHLIP and modify it with biotin. The modification appropriately increased the hydrophobicity and facilitated the formation of stable intramolecular hydrogen bonds of the peptide, both of which promoted the generation of the α-helical conformation. Thus, the resulting peptide, BAIP, could insert into the phospholipid bilayer for the isolation of EVs beyond the pH limitation. The simulation results and capture experiments confirmed the highly efficient (∼ 80% recovery rate) and unbiased enrichment of the EVs by the BAIP-based method, effectively overcoming the pH limitations.

Our BAIP-based isolation approach for EVs offers several advantages. (1) The meticulously-engineered peptide provided a size- and antigen-independent isolation approach, enabling the unbiased isolation of EVs. This would effectively address the challenge of missing EVs with low antigen expression due to the heterogeneity. (2) The BAIP-PS system can achieve a ∼ 80% recovery and significantly reduce the isolation time of EVs to 20 min. In comparison to existing methods, which often necessitate the isolation time ranging from 4 h to 22 h ^2, 21, 54^ and the capture efficiency from 20% to 80%, our BAIP-based method demonstrated significantly faster capture speed and more consistent capture efficiency. (3) The BAIP system exhibits exceptional anti-interference capability against non-target nucleic acids, proteins and other substances in plasma. The experiments confirmed that our BAIP-based method achieved a consistent capture efficiency of EVs in both buffer and clinical plasma samples. This capture efficiency is significantly higher than that of our previously-reported method, which utilized a lipid-nanoprobe but only achieved ∼ 50% capture efficiency in plasma samples^36^. (4) The purity of EVs by our method was better than that of the unbiased UC. In addition, our high quality and purified EVs can be used for downstream analysis, such as proteomics analysis and gene sequencing analysis. For the clinical trials, we successfully used the BAIP system in combination with qPCR to detect EGFR and KRAS gene mutations in EVs from clinical samples. (5) The versatility of BAIP allows for its conjugation with various bioinspired interfaces, enabling widespread applications in EVs isolation. (6) Our approach displayed a low cost of ∼ 7.5 $ to process each clinical sample (Supplementary Table 11), which was beneficial for the clinical translation. (7) Our BAIP system offers simplicity in operation and eliminates the need for expensive and bulky equipment, which may extend the potential applications in resource-limited areas.

Despite the encouraging findings of our study, there are some limitations and areas that can be improved in future research. Firstly, there are millions of possible reconfigurations available to modify the peptide, however, we have only investigated one specific reconfiguration. It is possible that other reconfigurations might prove to be more efficient in capturing the EVs in our ongoing studies. Anyhow, we have provided a new strategy for the peptide reconfiguration, particularly aiming at the EVs isolation, which could potentially be extended to modify the other peptides besides the WT pHLIP. Secondly, although our approach demonstrated some positive clinical trial outcomes, it is essential to conduct large-scale cohort studies to gather an extensive amount of clinical data. This will enable us to establish the relationship between the data and the corresponding cancer type or its origin, to extend the method for real clinical applications. Thirdly, the release of EVs from BAIP system can be achieved by simply cleaving the disulfide bonds with DTT or Tris (2-carboxyethyl) phosphine (TCEP). However, it is crucial to verify the biological function, biosafety, and the maintenance of structure integrity and bioactivity of the EVs for therapeutic purposes. The relevant research will be performed in the future. Fourthly, our unbiased approach for isolating and enriching EVs may play a vital role in identifying new biomarkers for cancer diagnostics using omics-based technology in clinical settings, holding great promise in enhancing the accuracy and effectiveness of EV-based cancer diagnosis^57^.

### Experimental Section

#### Chemicals and reagents

Dopamine, 2-(4-amidinophenyl)-6-indolecarbamidine dihydrochloride (DAPI) and albumin from bovine serum (BSA) were purchased from Aladdin (Shanghai, China). Sulfo-NHS-LC-Biotin and streptavidin (SA) were purchased from APExBIO (Houston, USA). Mouse anti-human antibodies against CD9 and Mouse anti-human antibodies against GAPDH were obtained from Abcam (Cambridge, UK). Cy3-labeled goat anti-mouse IgG was obtained from Boster Biological Technology co. Itd (Wuhan, China). SA-Cy3 and RNase-free H_2_O were obtained from Sangon Biotech Co., Ltd (Shanghai, China). The AIP was synthesized by Bankpeptide (Hefei, China). Ultrapure water was produced by Hitech-Kflow Water purification system (Shanghai, China).

#### Molecular dynamics simulations

We constructed the initial structures of peptides using the sequence of AIP in two forms, α-helix and loop, using PyMOL (Version 2.3, Schrödinger). Based on the structure of biotin used in the experiment, we used PyMOL to add the additional alkyl atoms to the original structure of biotin (obtained from the PDB: 1WPY). The all-atoms molecular dynamic simulations were performed using AMBER 16 package^58^. The AMBER ff14SB force field^59^ was used for the peptide, and the GAFF^60^ force field was used for the biotin. The initial structure was solvated in a cubic TIP3P water box with a 10 Å padding for all directions. A total of 40,000 steps minimization and 1ns equilibrate simulation were performed before the production process. The three independent simulation trajectories were performed for each system for 1μs at 298 K with a time step of 2 fs. The Particle Mesh Ewald (PME)^61^ was used to treat the long-range electrostatic interactions, and the short-range non-bonded interactions were calculated using a cutoff of 10 Å. The CPPTRAJ module was used for the analysis including in RMSD, RMSF, hydrogen bond and DSSP. The structure was rendered using PyMOL.

#### Functionalization of AIP with the biotin reagent

Briefly, 0.2 mg AIP and 0.29 mg Sulfo-NHS-LC-Biotin were dissolved in the mixed solution of PBS and DMSO (3:1, v/v) with shaking overnight. The solution was purified with a dialysis bag (1,000 Da) against PBS for 24 h with renewing PBS at 2 h, 4 h and 10 h. The obtained BAIP was kept in PBS containing 0.05% NaN_3_ and 1% BSA at −20 °C.

#### Verifying the binding of BAIP to cell membranes

The MCF-7 cells (human breast cancer cell line) were washed with PBS thrice and then incubated with BAIP for a certain time. After washed with PBS thrice and blocked with 5% BSA solution, the MCF-7 cells were incubated with SA-Cy3 for 15 min and then fixed with 4% paraformaldehyde for 10 min. After washed, the MCF-7 cells were stained with DAPI (1 μg mL^−1^) for 30 min. After thoroughly washed, the MCF-7 cells were imaged under an inverted fluorescence microscope (DSY5000X, China). Image J software was used to analyze the fluorescence intensities. For control group, the experiment was the same as above except that no BAIP was added. The influence of BAIP incubation time from 10 to 60 min was investigated and optimized.

#### Preparation and characterization of the BAIP-PS

Firstly, 1 mg dopamine was dissolved in 1 mL Tris-HCl buffer (10 mM, pH 8.5). Then 1 mg PS particles were homogeneously dispersed in the solution with shaking for 15 min. The obtained gray particles, named PS-PDA, were washed with ultrapure water for three times under 10,000 rpm for 5 min. Secondly, PS-PDA particles were homogeneously dispersed in 2 mL SA solution (10 mM Tris-HCl buffer, pH 8.5) with shaking for 30 min. The obtained particles, named PS-PDA-SA, were washed with PBS (10 mM, pH 7.4) for three times under 10,000 rpm for 5 min to remove unreacted SA. Thirdly, PS-PDA-SA particles were homogeneously dispersed in 1 mL BAIP solution (10 mM PBS, pH 7.4) with shaking for another 30 min. The obtained particles, named BAIP-PS, were washed as above to remove unreacted BAIP and kept in PBS containing 0.05% NaN_3_ and 1% BSA at 4°C. The morphology of these particles was examined by SEM (MIRA 3, TESCAN Brno, s.r.o., Russia). The zeta potential and size were measured using a Zetasizer.

#### Functionalized interfaces modified with BAIP

The ZnO-integrated microfluidic chip was fabricated according to our previous reports^62, 63^. The SA functionalized magnetic beads were purchased from MedChemExpress (shanghai, China). The PDA modified nickel foam was obtained as the previous reports^64^. All the functionalized interfaces modified with BAIP was similar to that of the PS.

#### Collection of clinical plasma samples

The blood samples of cancer patients and healthy person were obtained from Renmin Hospital of Wuhan University and Tongji Hospital of Huazhong University of Science and Technology (HUST). The study was reviewed and approved by the Institutional Review Board (IRB) of Tongji Hospital of HUST ([2021] IEC(A146). The whole blood samples were collected with 10 mL Vacutainer K2-EDTA tubes and immediately centrifuged at 300 g for 10 min at 4 [followed by centrifugation at 16,500 g for 20 min to remove microvesicles. The obtained plasma was filtered using a 0.22 μm filter and stored at −80 [for further use.

#### Cell culture

MCF-7 cells were cultured in Dulbecco’s modified Eagle’s medium (DMEM, Gibco) with 10% fetal bovine serum (FBS, Gibco) and 1% penicillin streptomycin (PS, Gibco). The non-small cell lung cancer cell lines A549 cells containing KRAS G12S mutations and PC9 cells containing EGFR exon 19 deletion mutations were cultured in RPMI-1640 medium with 10% FBS and 1% PS. All the cells were cultured in a 5% CO_2_ incubator at 37[.

#### Obtaining and characterization of model EVs

When 12 petri dishes in 10 cm diameter reached a confluency of 80%, the cells were cultured in the FBS-free medium for 48 h. The supernatant was collected and sequentially centrifuged at 4 [to remove cells and cellular debris (2000×g, 10 min), and microvesicles (10,000×g, 30 min), followed by filtration using a 0.22 μm filter. A total of 120 mL supernatant was thus collected and ultracentrifuged at a high speed of 120,000 g (2×80 min) at 4 °C using the type 70 Ti rotor. The obtained EVs pellets were re-suspended in 200 μL PBS as model EVs and stored at −80 [for further use.

For SEM, 10 μL of model EVs was fixed in 2.5% glutaraldehyde overnight and then dropped onto a poly-L-lysine-coated silicon wafer, subject to progressive dehydration with 20%, 30%, 50%, 70%, 85%, 95%, 100% gradient ethanol/H_2_O for 15 min per solution^65^. The sample was naturally dried and examined under a thermal field-emission environmental SEM.

For TEM, 10 μL of model EVs was dropped on a copper grid and adsorbed for 30 min at room temperature. Excessive EVs were sucked up with the filter paper and the EVs on the copper grid were negatively stained with 0.2% phosphotungstic acid for 1 min. Excessive phosphotungstic acid was absorbed with the filter paper. The sample was naturally dried at room temperature and then examined under a Talos F200X electron microscope (FEI, Netherlands).

For NTA, 10 μL of EVs was diluted with PBS to 1 mL and placed in the chamber. Nanoparticle Tracking Analysis software (NanoSight NS300, Malvern Instrument, England) was used to count the number of EVs.

#### Isolation and release of EVs

1 mg of the prepared BAIP-PS was added to 100 μL of EVs model sample with shaking at 500 rpm for a certain time at room temperature and then centrifuged at 10,000 rpm for 10 min. The supernatant was collected, and the RNA of the supernatant before and after capture was extracted. Qubit (Invitrogen, USA) was used to quantify the concentration of the extracted RNA. The capture efficiency (*Qc*) of EVs was calculated according to Eq. (1)

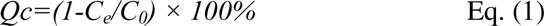

where *C_0_* and *C_e_* (ng/mL) are the initial and equilibrium concentrations of EVs RNA in solution, respectively. The influence of incubation time of BAIP-PS with EVs model samples from 10 to 40 min was assessed and optimized. The morphology of EVs-bound BAIP-PS was observed using SEM. Aliquots of 100 μL of plasma from healthy volunteer were shaken with 0.5, 1, 5, 10, 20 mg of the BAIP-PS, and the RNA was extracted as above to evaluate the capture efficiency.

The captured EVs were released with 100 μL of 100 μM DTT, and the released efficiency (*Qr*) of EVs was calculated according to Eq. (2)

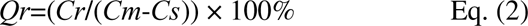

where *Cr* (ng/mL) was the concentration of released EVs RNA, *Cm* (ng/mL) and *Cs* (ng/mL) were the concentrations of model EVs RNA and supernatant EVs RNA, respectively.

#### Immuno-fluorescence staining

After capture of EVs, the BAIP-PS was thoroughly washed to remove non-specific adsorbed substances on the surface. Then, the EVs-bound BAIP-PS was incubated with mouse anti-human antibody against CD9 for 1.5 h at room temperature. After blocked with 5% BSA for 30 min, the particles were incubated with Cy3-labeled goat anti-mouse IgG for 30 min. Finally, the labelled BAIP-PS was observed using an inverted fluorescence microscope. For control group, the experiment was the same as above with BAIP-PS.

#### Nucleic acid and protein extraction

After isolation by BAIP-PS, the supernatant EVs were lysed with NucleoZol according to the manufacturer’s instructions to extract RNA. Typically, NucleoZol (500 μL) and RNase-free H_2_O (200 μL) were added into the supernatant with vigorously shaking for 15 s, then the mixed solution was incubated for 15 min at room temperature, and then centrifuged at 12,000 g for 15 min. The aqueous phase (500 μL) was homogenized with pure isopropanol (500 μL) and pelletized at room temperature for 10 min, followed by an RNA wash using 500 μL of 75% ethanol. Finally, the RNA pellet was dissolved in 20 μL of RNase-free water. The RNA concentrations were quantified using a Qubit Fluorometer. The mRNA reverse transcription PCR assays of CK19 gene and EGFR gene were performed as previously reported^66^.

The DNA from the isolated EVs was extracted using the QIAamp DNA micro kit (Qiagen, Germany) according to the manufacturer’s instructions. Briefly, after the EVs-bound BAIP-PS was thoroughly washed, lysis buffer (100 μL) and proteinase K (10 μL) were added to immerse the isolated EVs with heating at 56°C for 10 min. Afterwards, 50 μL of ethanol was added and the whole liquid phase was transferred to a spin column. The DNA was eluted in 30 μL of AE buffer after two washing steps and kept at −20°C until PCR amplification. The concentration of the extracted DNA was quantified with Nanodrop spectrophotometer (Thermo Scientific, USA). 2% agarose gel electrophoresis was used to identify the isolated DNA. The length of the fragments was determined using DNA ladders (1 kb).

To monitor GAPDH and CD9 expressed by EVs, the cells and EVs were separately lysed with ice-cold RIPA buffer containing protease inhibitors for 3.5 h. The lysates were centrifuged at 12,000 g for 10 min at 4 [and the proteins concentration was quantified using the BCA Kit (Sangon Biotech Co., Ltd, Shanghai). The proteins were further separated by SDS-PAGE and then transferred onto a polyvinylidene fluoride (PVDF) membrane. Subsequently, the PVDF membrane was blocked by 5% (w/v) skim milk in TBST buffer (2 mM Tris-HCl, 13.7 mM NaCl, and 0.1% Tween-20) at 37 [for 30 min. After washed with TBST buffer for three times, the membrane was incubated with primary antibody (1:1,000) overnight at 4[. After washed with TBST buffer for three times, the membrane was incubated with HRP-conjugated secondary antibodies for another 4 h. After washed three times with TBST buffer, the membrane was colored with BeyoECL Plus.

#### Sanger sequencing and qPCR assays

For Sanger sequencing, EGFR Exon 19 analysis of the isolated EVs DNA was performed as the previous report ^36^. The primers information was provided in Supplementary Table 12. The obtained first PCR products were cleaned using an AxyPrep PCR Clean-up Kit (Axygen, USA) following the manufacturer’s instructions. Then EGFR-mutant-enriched PCR assays were performed. Typically, 10-20 μL of the above PCR product was digested with Mse I at 37 °C overnight. An aliquot was employed as a template for the nest PCR amplification as the previous report ^36^. The obtained products were further purified and analyzed by Sanger sequencing (Applied Biosystems, USA). For qPCR assays, the DNA from the isolated EVs was measured with human EGFR and KRAS gene mutation detection kit (Yzymed, China) as previously reported ^66^.

#### Whole exome sequence

DNA whole-exome sequence was performed at the Department of Clinical Laboratory, Renmin Hospital of Wuhan University. The Ion AmpliSeq™ Library Kit 2.0 was used to prepare the sequencing library. WES was performed using an ion torrent platform (ThermoFisher, Shanghai, China). The data were quantitively assessed and filtered using Fastp^67^. Data were mapped to hg19 using bwa-mem (v.0.7.12)^68^ and coverage files were produced by Bedtools (v.2.17.0)^69^. IGV was used to visualize the mapping, and Bedtools was employed to calculate read counts within 10 kbp bins. Circus plots were generated to describe the read coverage in 10 kbp bins for each sample. The SNP and Indel of each sample were determined and filtered using GATK^70^, and the results of mutation were annotated using ANNOVAR^71^. The CNV of each sample was determine using control-FREEC^72^.

#### LC-MS/MS

Approximately 50 μg of proteins were denaturized at 37 [for 50 min with 10 mM DTT and alkylated with 40 mM iodoacetamide for 40 min in the dark, and then digested in 0.5 mL ultrafiltration units (Amicon Ultra-0.5, 30 kDa NMWL) with trypsin (1:50 w/w) in NH_4_HCO_3_ for 16 h at 37 °C. The obtained peptides were collected and dried by heating, which were then resuspended in 0.1% formic acid aqueous solution and analyzed using the Exactive^TM^ Plus Orbitrap high-resolution mass spectrometry and the EASY nLC^TM^1200 system (Thermo Fisher Scientific). A C18 column (15 cm, 75 μm id, 3 μm, 100 Å) with a flow rate of 600 nL /min was used for peptides separation. The RAW MS files were processed using the Proteome Discoverer v2.1 (Thermo Fisher Scientific, Rockford, USA) against the human protein database from OpenProt (https://openprot.org/). The parameters were set up as follows. Enzyme: trypsin (full) with maximum missed cleavages was 2; precursor mass tolerance was set to 10 ppm; fragment mass tolerance: 0.02 Da; false discovery rate was defined as 1%. All comparisons with references were obtained by downloading the proteomics raw data of reference and then conducting relevant analyses.

#### Statistical analysis

Data were expressed as mean ± SD. Statistical analyses were assessed by two-tailed t test with significance at *P* < 0.05. Analyses were performed with Origin 9.0.

## Supporting information

Supplementary data associated with this article can be found in the online version at doi:xxxxxxxxxxxxxxxx.

## Author Contributions

L. W. designed the research, conducted experiments, analyzed data and wrote and edited original draft. Z. G. assisted with molecular dynamics simulations. M. W. designed the research, assisted with the analysis of WES data and provided the clinical samples. Y.-Z. L. assisted with the measurement of the hydrated particle size and zeta potential of materials. J. Z. provided the clinical samples. Q. X. assisted with the analysis of WES data. M. J. wrote the original draft. X.-W. W. discussed the results. Q.-Y. L. edited the manuscript. C. Z. discussed the results. L.-Y. M. edited the manuscript. S.-Y. Z. discussed the results. X. Y. provided technical and advisory, supervised the project. L. X. provided technical and advisory, supervised the project. All authors have given approval to the final version of the manuscript.

## Acknowledgements

This work is supported by the National Natural Science Foundation of China (Grant No. 21804105, 22174049, 31971155), the Program for HUST Academic Frontier Youth Team (Grant No. 2019QYTD09), the Natural Science Foundation of Hubei Province of China (No.2021CFB335), the Fundamental Research Funds for the Central Universities in China (Grant Nos. 2172019KFYRCPY112, 2172020kfyXJJS082) and the Youth Innovation Promotion Association of the Chinese Academy of Sciences (Grant No. 2020329). The authors thank the Analytical and Testing Center of HUST for the material characterization.

